# Merging Bioactivity Predictions from Cell Morphology and Chemical Fingerprint Models Using Similarity to Training Data

**DOI:** 10.1101/2022.08.11.503624

**Authors:** Srijit Seal, Hongbin Yang, Maria-Anna Trapotsi, Satvik Singh, Jordi Carreras-Puigvert, Ola Spjuth, Andreas Bender

## Abstract

The applicability domain of machine learning models trained on structural fingerprints for the prediction of biological endpoints is often limited by the lack of diversity of chemical space of the training data. In this work, we developed similarity-based merger models which combined the outputs of individual models trained on cell morphology (based on Cell Painting) and chemical structure (based on chemical fingerprints) and the structural and morphological similarities of the compounds in the test dataset to compounds in the training dataset. We applied these similarity-based merger models using logistic regression models on the predictions and similarities as features and predicted assay hit calls of 177 assays from ChEMBL, PubChem and the Broad Institute (where the required Cell Painting annotations were available). We found that the similarity-based merger models outperformed other models with an additional 20% assays (79 out of 177 assays) with an AUC>0.70 compared with 65 out of 177 assays using structural models and 50 out of 177 assays using Cell Painting models. Our results demonstrated that similarity-based merger models combining structure and cell morphology models can more accurately predict a wide range of biological assay outcomes and further expanded the applicability domain by better extrapolating to new structural and morphology spaces.

Figure:
For TOC Only

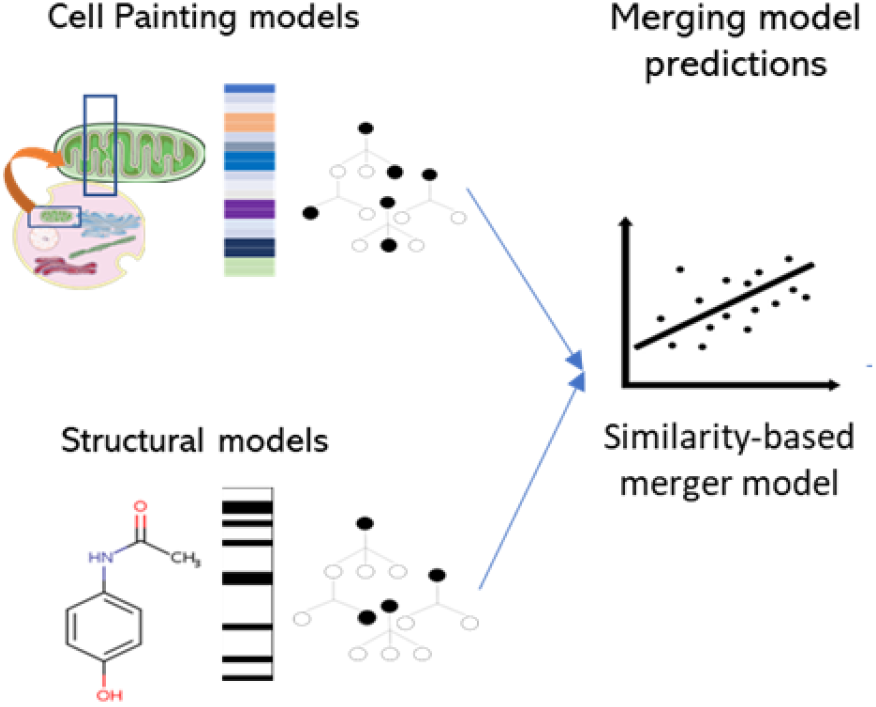

## INTRODUCTION

The prediction of bioactivity, mechanism of action (MOA)^1^, safety and toxicity^2^ of compounds using only chemical structure is challenging given that such models are limited by the diversity in the chemical space of the training data.^3^ The chemical space of this data on which the model is trained is used to define the applicability domain of the model.^4^ Among the various ways to calculate a model’s applicability domain, Tanimoto similarity for chemical structure is commonly used as a benchmark similarity measure for compounds. Tanimoto distance-based Boolean applicability has been previously used to improve the performance of classification models.^5^ Expanding the applicability domain of structural models will improve the reliability of a model to predict endpoints for new compounds. One way to achieve this would be to incorporate hypothesis-free high-throughput data, such as cell morphology^6^, bioactivity data^7^ or predicted bioactivities^8,9^ in addition to structural models.^10^ This then has the potential to improve predictions for compounds structurally distant from the training data. This is because compounds having similar biological activity may not always have a similar structure; however, they may show similarities in the biological response space.^11^ Recently, using Chemical Checker signatures derived from processed, harmonized and integrated bioactivity data, researchers demonstrated that similarity extends well beyond chemical properties into biological activity throughout the drug discovery pipeline (from *in vitro* experiments to clinical trials).^12^ Hence the use of biological data could significantly help predictive models that have often been trained solely on chemical structure.^10^

In recent years, relatively standardized hypothesis-free cell morphology data can now be obtained from the Cell Painting assay.^13^ Cell Painting is a cell-based assay that, after a given chemical or genetic perturbation, uses six fluorescent dyes to capture a snapshot of the cellular morphological changes induced by the aforementioned perturbation. The six fluorescent dyes allow for the visualization of eight cellular organelles, which are imaged in five-channel microscopic images. The microscopic images are typically further processed using image analysis software, such as Cell Profiler^14^, which results in a set of circa 1700 morphological numerical features per cell. These numerical features representing morphological properties such as shape, size, area, intensity, granularity, and correlation, among many others, are considered versatile biological descriptors of a system.^6^ Previous studies have shown Cell Painting data to be predictive of a wide range of bioactivity and drug safety-related endpoints such as the mechanism of action^15^, cytotoxicity^16^, microtubule-binding activity^17^, and mitochondrial toxicity^18^. Recently, it has also been used to identify phenotypic signatures of PROteolysis TArgeting Chimeras (PROTACs)^19^ as well as to determine the impact of lung cancer variants^20^. Thus, Cell Painting data can be expected to contain a signal about the biological activity of the compound perturbation,^6^ and in this work, we explored how best to combine Cell Painting and chemical structural models for the prediction of a wide range of biological assay outcomes.

From the modeling perspective, several ensemble modeling techniques have been proposed to combine predictions from individual models.^21^ One way to achieve this is an ensembling method shown in Figure 1a, referred to as a soft-voting ensemble in this work. This method computes the mean of predicted probabilities from individual models and thus provides equal weight to individual model predictions. However, soft-voting ensemble models when combining two individual models give equal importance to each model.^21^ This implies that if a model predicts a higher probability for a compound to be active and another model predicts the same compound to be inactive but with a lower probability, the first model prediction is considered final without considering the individual model’s reliability. As shown in Figure 1b, another way to combine predictions from different models is *via* model stacking where the predictions of the individual models are used as features to build a second-level model (referred to as a hierarchical model in this work). Hierarchical models have previously been used by integrating classification and regression tasks in predicting acute oral systematic toxicity in rats.^22^ The applicability range of predictions can be estimated by (i) the Random Forest predicted class estimates^23^ (referred to as predicted probabilities in this study) and (ii) using the similarity of the test compound to training compounds (which in turn approximates the reliability of the prediction).^24^ The hypothesis of the current work is hence that using the similarity of the test compound to training compounds in individual feature spaces and predicted probabilities of individual models built on those feature spaces can improve the model performance.

**Figure 1:**
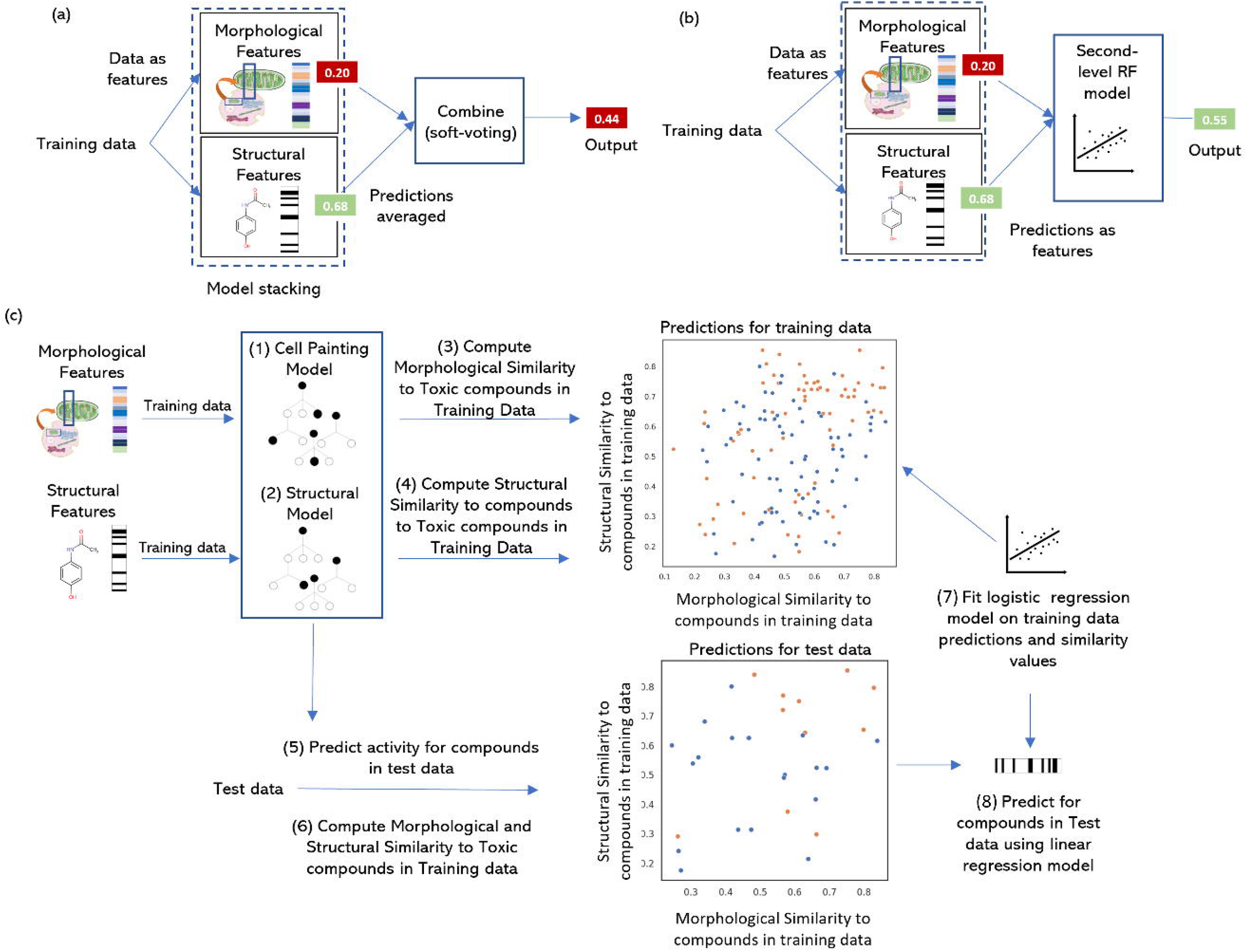
Schematic Representation of workflow in this study to build (a) hierarchical models where the predictions of the individual models are used as features to build a second-level model, and (b) soft-voting ensemble models that compute the mean of predicted probabilities from individual models and (c) the similarity-based merger model. The similarity-based merger model combined predicted probabilities from individual models and the morphology and structural similarity of compounds to active compounds in training data.

The various ways of fusing structural models with models trained on cell morphology were recently exploited by Moshkov et al.^27^ who used chemical structures and cell morphology data (from the Cell Painting assay) to predict the compound activity of 270 anonymised bioactivity assays from academic screenings in the Broad Institute. They used a late data fusion (by using a majority rule on the prediction scores similar to soft-voting ensembles) to merge predictions for individual models. The late data fusion models were able to predict 31 out of 270 assays with AUC>0.9, compared with 16 out of 270 assays for models using only structural features. This showed that fusing models built on two different feature spaces that provide complementary information were able to improve the prediction of bioactivity endpoints. Previous work has also shown that combinations of descriptors can significantly improve prediction for MOA classification^25,26,15^ (using gene expression and cell morphology data), cytotoxicity^16^, mitochondria toxicity^18^ and anonymised assay activity^27^ (using chemical, gene expression, cell morphology and predicted bioactivity data), prediction of sigma 1 (σ1) receptor antagonist^28^ (using cell morphology data and thermal proteome profiling), and even organism-level toxicity^29^ (using chemical, protein target and cytotoxicity qHTS data). Thus, the combination of models built from complementary feature spaces can expand a model’s applicability domain by allowing predictions in new structural space.^30^

In this work, we explored merging predictions of assay hit calls from chemical structural models with predictions from another model using Cell Painting data for 88 assays from public datasets from PubChem and ChEMBL (henceforth referred to as public dataset, assay descriptions released as Supplementary Data 1) and 89 anonymised assays from the Broad Institute^27^ (henceforth referred to as Broad Institute dataset, assay descriptions released as Supplementary Data 2). Cell Painting data, in general, may be assumed to be only highly predictive of the cell-based assay. However, in this study, we did not specifically select assays where this relation was obvious, as that would make our comparisons significantly favour the Cell Painting assay. In this work, we simply compare the two feature spaces, and for this, we use a wide range of assays (as mentioned above) while also later interpreting which feature spaces work better for which particular assays. That being noted, the Cell Painting assay is being constantly investigated for signals in not just *in vitro* assays but also *in vivo* effects; recent studies have established a signal for lung cancer^20^ and drug polypharmacology^31^.

From the modelling perspective, as shown in Figure 1c, we merged predictions using a logistic regression model that not only takes the predicted probabilities from individual models but also the test compound’s similarity to the active compounds in the training data in different feature spaces. That is, the models are also provided with the knowledge of how morphologically/structurally similar the test compound is to other active compounds in the training set. Here we emphasise using similarity-based merger models to improve the applicability domain of individual models (predicting compounds that are distant to training data in respective feature spaces) and the ability to predict a wider range of assays with the combined knowledge from the chemical structure and biological descriptors from Cell Painting assay.

## RESULTS AND DISCUSSIONS

The 177 assays used in this study are a combination of the public dataset and anonymised assays from a Broad Institute dataset where required Cell Painting annotations were available (see Methods section for details). The public dataset comprising 88 assays (with at least 100 compounds) was collected from Hofmarcher et al^38^ and Vollmers et al^40^ (see Supplementary Data 1 for assay descriptions) for which Cell Painting annotations were available from the Cell Painting assay^46^. The Broad Institute dataset comprises 89 assays (as shown in Supplementary Data 2 for assay descriptions). We trained individual Cell Painting and structural models for all 177 assays. We used two baseline models for comparison, namely a soft-voting ensemble and a hierarchical model. Finally, we compared the results from the individual models and baseline ensemble models to the similarity-based merger models.

### Similarity-based merger models outperform other baseline models

As shown in Figure 2, we found that similarity-based merger models performed with significantly improved AUC-ROC (mean AUC 0.66 using similarity-based merger models,) compared with Cell Painting models (mean AUC 0.62 using, p-value from paired t-test of 5.6×10^-4^) and structural models (mean AUC 0.64, p-value from paired t-test of 7.3×10^-3^) for 171 out of the 177 assays (all models for the remaining 6 assays have AUC<0.50, hence any improvement is insignificant as the models’ performance remains worse than random). Figure S1 shows that similarity-based merger models significantly improved Balanced Accuracy and F1 scores compared with individual models. Overall, similarity-based merger models outperform other models in predicting bioactivity endpoints.

**Figure 2:**
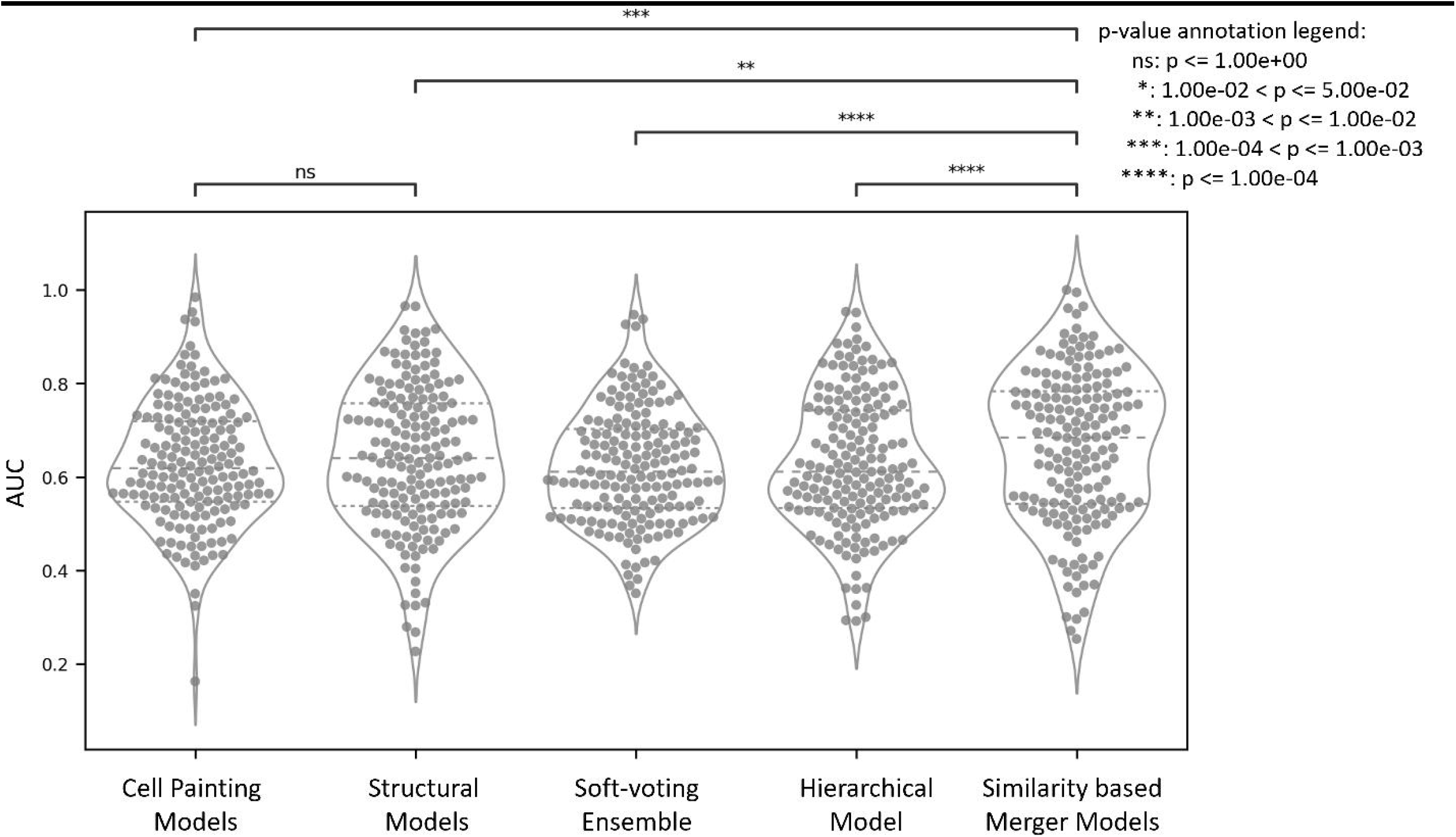
Distribution of AUC of all models, Cell Painting, Morgan Fingerprints, baseline models of a soft-voting ensemble, hierarchical model, and the similarity-based merger model, over 171 assays (out of 177 assays). An assay was considered for a paired significance test only if the balanced accuracy was >0.50 and the F1 score was >0.0 for at least one of the models.

As shown in Figure 3, 79 out of 177 assays achieved AUC>0.70 with the similarity-based merger model, followed by hierarchical models for 55 out of 177 assays. Structural models achieved AUC>0.70 in 65 out of 177 assays while for the Cell Painting models, this was the case in 50 out of 177 assays. Further 25 assays out of 177 were predicted with AUC>0.70 with all methods while only 12 out of 177 assays did not achieve AUC>0.70 with similarity-based merger models but did with the other models. When considering balanced accuracy, 51 out of 177 assays achieved a balanced accuracy > 0.70 with similarity-based merger models compared with 44 out of 177 assays for soft-voting ensemble models, as shown in Figure S2.

**Figure 3:**
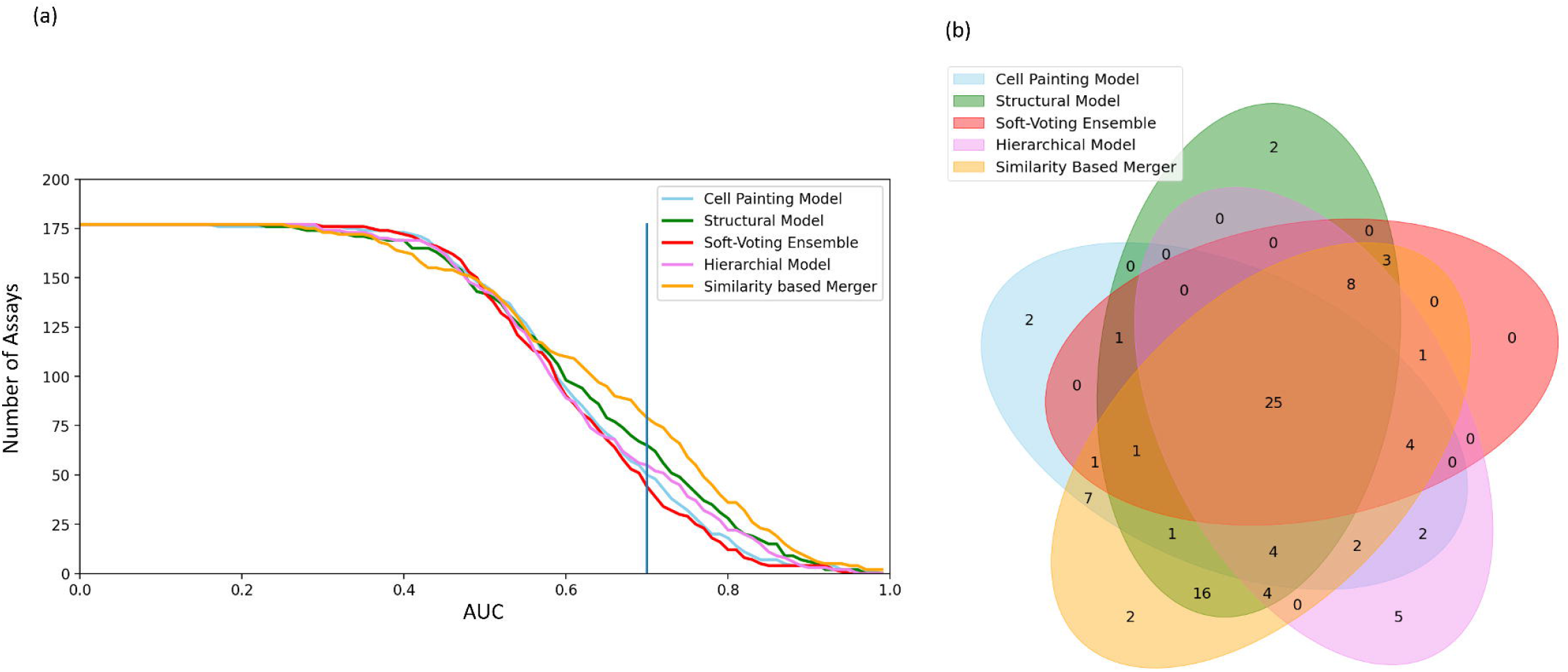
(a) Number of assays predicted with an AUC above a given threshold. (b) Distribution of assays with AUC > 0.70 common and unique to all models, Cell Painting, Morgan Fingerprint, baseline models of a soft-voting ensemble, hierarchical model, and the similarity-based merger model, over all of the 177 assays used in this study.

Comparing performance for the Cell Painting and structural models by AUC individually (Figure S3) we observed that structural models and Cell Painting models were complementary in their predictive performance; while 96 out of 177 assays achieve a higher AUC with structural information alone, 81 out of 177 assays achieve a higher AUC using morphology alone as shown in Figure S3a. Hierarchical models outperform soft-voting ensembles for 106 out of 177 assays as shown in Figure S3b. Finally, the similarity-based merger model achieved a higher AUC score for 124 out of 177 assays compared with 52 out of 177 with hierarchical models and 132 out of 177 assays compared with 45 out of 177 with soft-voting ensembles as shown in Figures S3c and S3d. This shows that the similarity-based merger model was able to leverage information from both Cell Painting and structural models to achieve better predictions in assays where no individual models were found to be predictive thus indicating a synergistic effect.

We next looked at the performance at the individual assay level (as shown in Supplementary Data 3) as indicated by the AUC scores. We looked at 162 out of 177 assays where either the similarity-based merger model or the soft-voting ensemble performed better than a random classifier (*AUC* = 0.50) We observed that for 127 out of 177 assays (individual changes in a performance recorded in Figure S4), the similarity-based merger models improved performance compared with the soft-voting ensemble (with the largest improvement recorded at 65.1%) and a decrease in performance was recorded in 35 out of 177 assays (largest decrease recorded at −58.0% in performance). Further comparisons of AUC performance in Figure S5 show that similarity-based merger models improved AUC compared with both structural models and Cell Painting models. This improvement in AUC was independent of the total number of compounds in the assays as shown in Figure S6. Thus, we conclude that the similarity-based merger model outperformed individual models by combining the rich information contained in cell morphology and structure-based models more efficiently than baseline models.

### Similarity-based merger models expand the applicability domain compared with individual models

We next determined how individual and similarity-based merger model predictions differ with compounds that were structurally or morphologically similar/dissimilar to active compounds in the training set. We looked at predictions for each compound from the Cell Painting and structural models over the 177 assays and grouped them based on their morphological and structural similarity to active compounds in the raining set respectively. We observed, as shown in Figure S7, that similarity-based merger models correctly classified a higher proportion of test compounds which were less similar morphologically to active compounds in the training data. Further, as the structural similarity of test compounds with respect to active compounds in the training set increased, the structural models correctly classified a higher proportion of compounds while similarity-based merger models correctly classified test compounds with both low and high structural similarity. For example, out of 360 compounds with a low structural similarity between 0.20 to 0.30, models using chemical structure correctly classified 56.2% of compounds while similarity-based merger models correctly classified a much greater 63.6% of compounds. At the same time, out of 1,525 compounds with higher structural similarity between 0.90 to 1.00, models using chemical structure correctly classified 75.5% of compounds compared with the similarity-based merger models that correctly classified 75.2% of compounds. This shows that the similarity-based merger model correctly predicted a larger proportion of compounds over a wide range of structural and morphological similarities to the training set, hence demonstrating an increase in the applicability domain.

For clarity of the reader, this is further illustrated in Figure S8 as in the case of a particular assay, namely 240_714 from the Broad Institute, a fluorescence-based biochemical assay. Here, the structural model correctly predicted toxic compound activity when they were structurally similar to the training set. The Cell Painting model performed better over a wide range of structural similarities but was often limited when morphological similarity was low. The similarity-based merger models learned and adapted across individual models from local regions in this structural versus morphological similarity space in a manner best suited to compounds in that region to correctly classify a wider range of active compounds with lowered structural and morphological similarities to the training set.

### Comparison of Performance at Gene Ontology Enrichment level

We next analysed the assays (and associated biological processes) where the Cell Painting model, the structural model, and the similarity-based merger model were most predictive and therefore if there was complementary information present in both feature spaces. Results presented here are from the PubChem dataset comprising 88 assays as the Broad Institute dataset is not annotated with complete biological metadata, which renders some of the more detailed analysis downstream not viable.

Figure 4a shows a protein-protein network (annotated by genes) from the STRING database labelled by the model performance where the respective individual model was better predictive (or otherwise equally predictive, which includes cases where different models are better predictive over multiple assays related to the same protein target). We found meaningful models (AUC > 0.50) were achieved for 27 out of 34 gene annotations when using the Cell Painting and for 25 out of 34 gene annotations using the structural model. Of these, the Cell Painting models were better predictive for 25 out of 32 gene annotations (mean AUC= 0.65) compared with the structural models which were better predictive for 23 out of 32 gene annotations (mean AUC=0.56). We next compared the hierarchical model to the similarity-based merger model for 35 gene annotations where either model achieved AUC>0.50. The hierarchical model performed with higher AUC (mean AUC= 0.57) for only 4 out of 34 gene annotations compared with the similarity-based merger model which was better predictive for 23 out of 34 gene annotations (mean AUC=0.60). Thus, we observed that similarity-based merger models performed better over a range of assays (over 23 out of 34 gene annotations) capturing a wide range of biological pathways.

**Figure 4:**
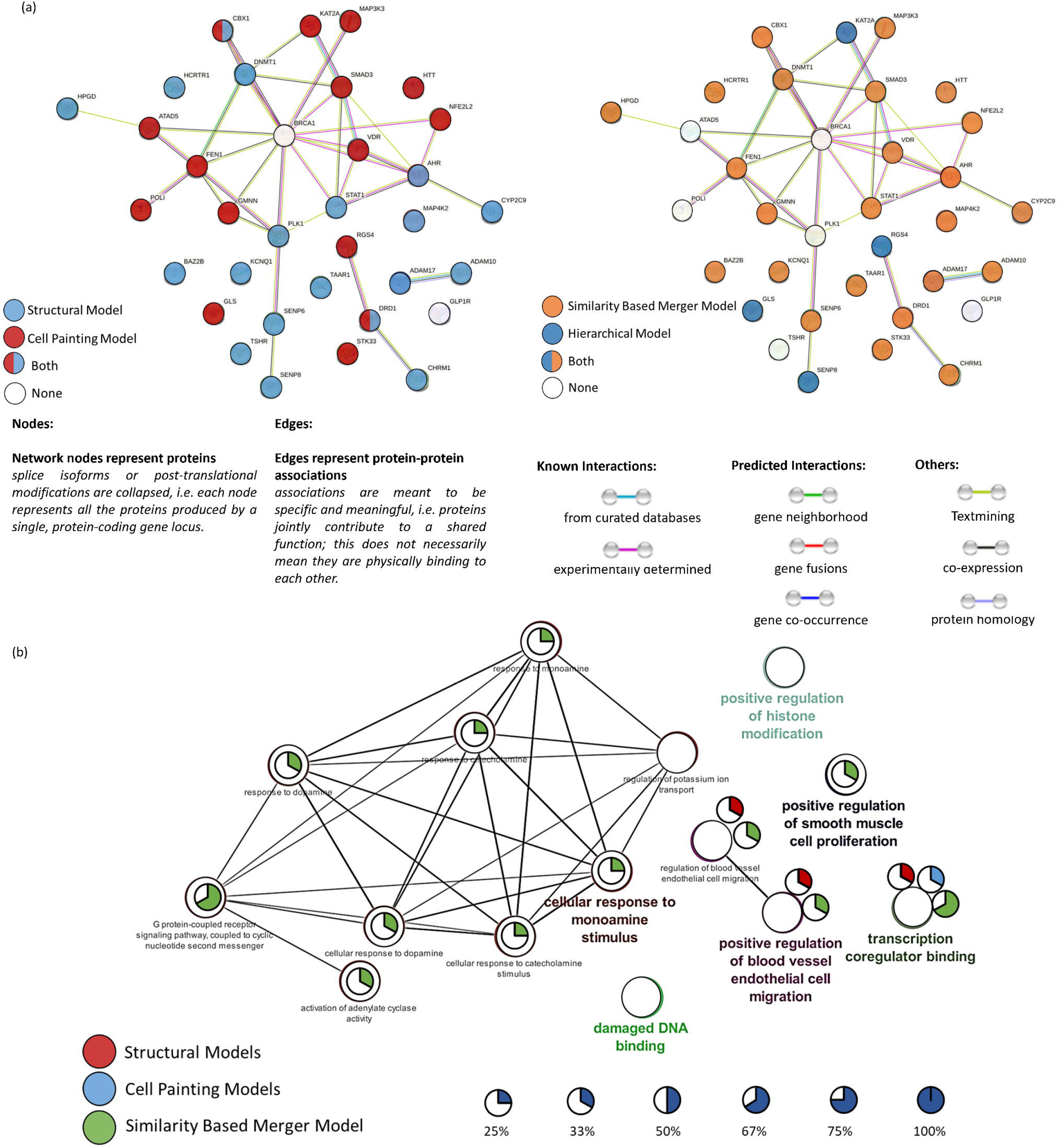
(a) STRING gene-gene interaction networks for 34 Genes annotations associated with 37 assays in the public dataset labelled by the model which was better predictive compared with the other models and a random classifier with an AUC>0.50 (b) Molecular and functional pathway terms related to the 37 assays using the Cytoscape^43^ v3.9.1 plugin ClueGO^44^ labelled by percentage of gene annotations where an AUC>0.70 was achieved by the Cell Painting, structural and similarity-based merger models.

Cell Painting models performed better than structural models for assays associated with 6 gene annotations: ATAD5, FEN1, GMNN, POLI and STK33 (with an average AUC = 0.64 for Cell Painting models compared with AUC = 0.48 for structural models). These gene annotations were associated with molecular functions of ‘GO:0033260 Nuclear DNA replication’ and ‘GO:0006260 DNA replication’ which are processes resulting in morphological changes, which were captured by Cell Painting. Among gene annotations associated with the assays better predicted by structural models are TSHR, TAAR1, HCRTR1 and CHRM1 (with an average AUC = 0.70 for structural models compared with an AUC = 0.63 for Cell Painting models). These gene annotations are associated with the KEGG pathway of ‘neuroactive ligand-receptor interaction’ and the Reactome pathway of ‘amine ligand-binding receptors’ which were captured better by chemical structure. Hence, we see that Cell Painting models perform better on assays capturing morphological changes in cells or cellular compartments such as the nucleus, while structural models work better for assays associated with a ligand-receptor activity. In addition, the KEGG term ‘amine ligand-binding receptors’ is defined on the chemical ligand level explicitly, making the classification of compounds falling into this category from the structural side easier. The similarity-based merger models hence combined the power of both spaces and were predictive for assays affecting morphological changes (average AUC= 0.58 for the similarity-based merger model) as well as related to the ligand-receptor binding activity (average AUC= 0.78 for similarity-based merger model).

This is further illustrated in Figure 4b which shows enriched molecular and functional pathway terms from ClueGO^44^ for the 34 gene annotations available. Both Cell Painting models and structure-based models were limited to predicting with AUC>0.70 only 33% of gene annotations associated with only two pathways, namely, transcription coregulator binding and positive regulation of blood vessel endothelial cell migration pathways. On the other hand, similarity-based merger models predicted 25-67% gene annotations associated with multiple pathways with an AUC>0.70. These pathways included transcription coregulator binding, positive regulation of blood vessel endothelial cell migration pathways, positive regulation of smooth muscle cell proliferation and G protein-coupled receptor signalling pathways among others. Hence this underlines the utility of similarity-merger models across a range of biological endpoints.

### Comparison of Performance by Readout and Assay type

Results presented here are from the Broad Institute dataset comprising 89 assays (as shown in Supplementary Data 3) which were released with only information about assay type and readout type (for details see Supplementary Data 2 and Figure S9); we analysed the Cell Painting, structural and similarity-based merger model as a function of those.

As shown in Figure 5, Cell Painting models perform significantly better with a relative 8.8% increase in AUC with assays measuring luminescence (mean AUC = 0.72) compared with assays measuring fluorescence (mean AUC = 0.66) while structural and similarity-based merger model show no significant differences in performances. The better predictions in the case of luminescence-based assays, which are readouts specifically designed to answer a biological question, and can be related to the use of a reporter cell line and a reagent that based on the ATP content of the cell, is converted to a luciferase substrate which leads to a cleaner datapoint. ^32^ On the other hand, Cell Painting is an unbiased high-content imaging assay that takes into consideration the inherent heterogeneity in cell cultures where we visualise cells (often even measuring at a single cell level), contrary to a luminescence assay where one measures the average signal of a cell population. Further Cell Painting models performed significantly better with a relative 18.1% increase in AUC for cell-based assays (mean AUC = 0.72) compared with biochemical assays (mean AUC = 0.61). This might be due to also the Cell Painting assay being a cellular assay, hence also implicitly including factors such as membrane permeability in measurements. Further, most similarity-based merger models outperform baseline models over assay and readout types as shown in Figures S10 and S11. Overall, Cell Painting models can hence be considered to provide complementary information to chemical structure regarding cell-based assays, which was particularly beneficial for the significant improvement in the performance of similarity-based merger models.

**Figure 5:**
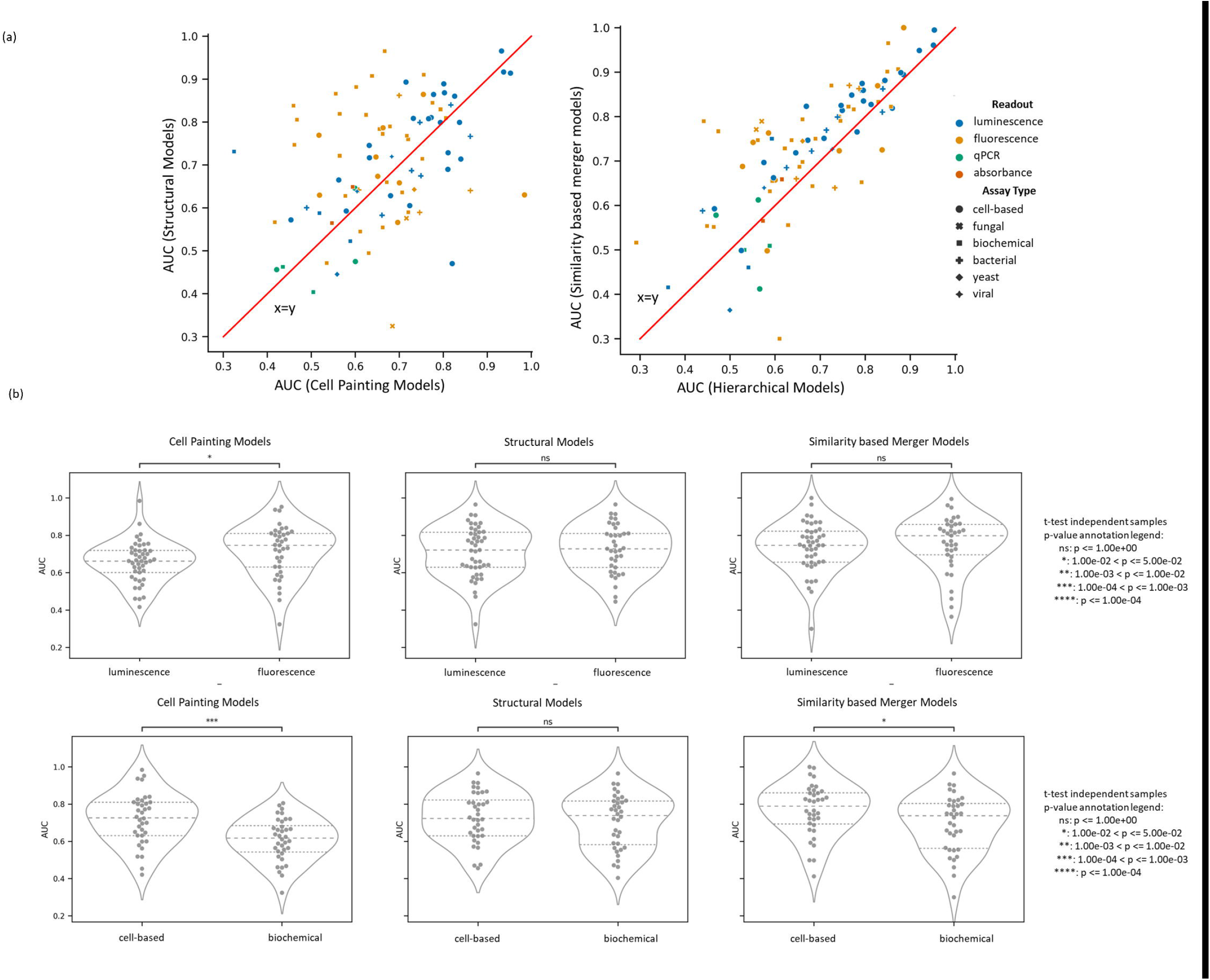
AUC performance of models using Cell Painting, structural models, and similarity-based merger model for 89 assays in the Broad Institute dataset based on readout type (fluorescence and luminescence) or the assay type (cell-based and biochemical). Further details are shown in Figures S10 and S11.

### Limitations of this work

One limitation of the study design is having to balance unequal data classes by under-sampling. Here, the data was therefore initially under-sampled to a 1:3 ratio of majority to minority class to build a similarity-based merger model, which leads to some loss of experimental data. Further, after splitting the dataset into training and test datasets, the training data needs to contain enough samples spread across the structural versus morphological similarity map for the models to work well. This was ensured by a random split; other splitting strategies such as scaffold-based splitting may not allow the use of the second-level models as the chemical/biological space of the test data will vary significantly from the training data. Finally, the current study design is also affected by methodological limitations.^33,34^ In the current study design, the explicit definition of similarity of a compound in chemical and morphological space, which although used here for better interpretability, could have been possible via different ways of learning the data directly, for example using Bayesian inference.^35^

From the side of feature spaces, Cell Painting data is derived from U2OS cell-based assays which are usually different from the cell lines used in measuring the activity endpoint. However previous work has shown that Cell Painting data is similar across different cell lines and the versatile information present was universal, that is, the genetic background of the reporter cell line does not affect the AUC values for MOA prediction.^36^ Thus Cell Painting data can be used to model different assays with different cell lines. Future studies will also benefit from larger datasets, such as the JUMP-CP consortium.^37^

## CONCLUSIONS

Predictive models that use chemical structures as features are often limited in their applicability domain to compounds which are structurally similar to the training data. To the best of our knowledge, this is the first paper which uses both similarity and predictions from chemical structural and cell morphology feature spaces to predict assay activity. Our results should have clear implications for similarity-based merger models (that are shown to be comparatively better than baseline soft-voting ensembles and hierarchical models) and can be used to predict bioactivity over a wide range of small compounds. In this work, we developed similarity-based merger models to combine two models built on complementary feature spaces of Cell Painting and chemical structure and predicted assay hit calls from 177 assays (88 assays from the public dataset and 89 assays from a dataset released by the Broad Institute) for which Cell Painting data were available.

We found that Cell Painting and chemical structure contain complementary information and can predict assays associated with different biological pathways, assay types, and readout types. Cell Painting models achieved higher AUC better for cell-based assays and assays related to biological pathways such as DNA replication. Structural models achieved a higher AUC for biochemical and ligand-receptor interaction assays. The similarity-based merger models, combining information from the two feature spaces, achieved a higher AUC for cell-based (mean AUC=0.77) and biochemical assays (mean AUC=0.70) as well as assays related to both biological pathways (mean AUC=0.58) and ligand-receptor based pathways (mean AUC=0.74). Further, the similarity-based merger models outperformed all other models with an additional 20% assays with AUC>0.70 (79 out of 177 assays compared with 65 out of 177 assays using structural models). We also showed that the similarity-based merger models correctly predicted a larger proportion of compounds which are comparatively less structurally and morphologically similar to the training data compared with the individual models, thus being able to improve the applicability domain of the models.

In conclusion, the similarity-based merger models greatly improved the prediction of assay outcomes by combining high predictivity of fingerprints in areas of structural space close to the training set with better generalizability of cell morphology descriptors at greater distances to the training set.

On the practical side, Cell Painting assay is a single screen-based hypothesis-free assay that is inexpensive compared with dedicated assays. Being able to use such an assay for bioactivity prediction will greatly improve the cost-effectiveness of such assays. Similarity-based merger models used in this study can hence improve the performance of predictive models, particularly in areas of novel structural space thus contributing to overcoming the limitation of chemical space in drug discovery projects.

## METHODS

### Bioactivity Datasets

We retrieved drug bioactivity data as binary assay hit calls for 202 assays and 10,570 compounds from Hofmarcher et al^38^ who searched ChEMBL^39^ for assays for which cell morphology annotations from the Cell Painting assay were available as shown in Figure S12. We further added binary assay hit calls from another 30 assays not included in the source above from Vollmers et al^40^ who searched PubChem^41^ assays for overlap with Cell Painting annotations. Additionally, we used 270 anonymised assays (with binary endpoints) from the Broad Institute^27^ as shown in Figure S12b. This dataset, although not annotated in with biological metadata, comprises assay screenings performed over 10 years at the Broad Institute and is representative of their academic screenings.

### Gene Ontology Enrichment of Bioactivity Assays

From the public dataset of 88 assays used in this study where detailed assay data was available, 37 out of 88 assays where experiments used human-derived cell lines were annotated to 34 protein targets. Next, we determined using the STRING database^42^, we annotated all 34 protein targets with the associated gene set and further obtained a set of Gene Ontology terms associated with the protein target. We used Cytoscape^43^ v3.9.1 plugin ClueGO^44^ to condense the protein target set by grouping them into functional groups to obtain the associated significance (using the baseline ClueGO p-value ≤0.05) molecular and functional pathway terms. In this manner, we associated individual assays with molecular and functional pathways for further evaluation of model performances.

### Cell Painting Data

The Cell Painting assay used in this study, from the Broad Institute, contains cellular morphological profiles of more than 30,000 small molecule perturbations.^45,46^ The morphological profiles in this dataset are composed of a wide range of feature measurements (share, area, size, correlation, texture etc. as shown in a demonstrative table in Figure S12a). While preparing this dataset, the Broad Institute normalized morphological features to compensate for variations across plates and further excluded features having a zero median absolute deviation (MAD) for all reference cells in any plate.^13^ Following the procedure from Lapins et al, we subtracted the average feature value of the neutral DMSO control from the particular compound perturbation average feature value on a plate-by-plate basis.^15^ For each compound and drug combination, we calculated a median feature value. Where the same compound was replicated for different doses, we used the median feature value across all doses that were within one standard deviation of the mean dose. Finally, after SMILES standardisation and removing duplicate compounds using standard InChI calculated using RDKit^48^, we obtained 1783 median Cell Painting features for 30,404 unique compounds (available on Zenodo at https://doi.org/10.5281/zenodo.7589312).

### Overlap of Datasets

For both the public and Broad dataset, as shown in Figure S12b (step 1) we used MolVS^47^ standardizer based on RDKit^48^ to standardize and canonicalize SMILES for each compound which encompassed sanitization, normalisation, greatest fragment chooser, charge neutralisation, and tautomer enumeration described in the MolVS documentation^47^. We further removed duplicate compounds using standardised InChI calculated using RDKit^48^.

Next, for the public dataset, we obtained the overlap with the Cell Painting dataset using standardised InChI Figure S12b (step 2). From this, we removed assays which contained less than 100 compounds for the minority class with Cell Painting datasets (which were difficult to model due to limited data) as shown in Figure S12b (step 3). For each assay, as shown in Figure S12b (step 4)the majority class (most often the negative class) was randomly resampled to maintain a minimum 3:1 ratio with the minority class to ensure that models are fairly balanced. Finally, we obtained the public assay data for a sparse matrix of 88 assays and 9,876 unique compounds (see Supplementary Data 1 for assay descriptions). Similarly, for the Broad dataset, out of 270 assays provided, as shown in Figure S12b, we removed assays that contained less than 100 compounds and randomly resampled to maintain a minimum 3:1 ratio with the minority class, resulting in a Broad Institute dataset as a sparse matrix of 15,272 unique compounds over 89 assays (see Supplementary Data 2 for assay descriptions). Figure S13 shows the distribution of the total number of compounds for 177 assays used in this study. Both datasets are publicly available on Zenodo at https://doi.org/10.5281/zenodo.7589312).

### Structural Data

We generated Morgan Fingerprints of radius 2 and 2048 bits using RDKit^48^ used as binary chemical fingerprints in this study (as shown in a demonstrative table in Figure S12a).

### Feature Selection

Firstly, we performed feature selection to obtain morphological features for each compound. From 1,783 Cell Painting features, we removed 55 blocklist features that were known to be noise from Way et al.^49^ For the compounds in the public assays, we further removed 1,012 features which had a very low variance below a 0.005 threshold using the scikit-learn^50^ variance threshold module. Next, similar to the feature section implemented in pycytominer^51^, we obtained the list of features such that no two features correlate greater than a 0.9 Pearson correlation threshold. For this, we calculated all pairwise correlations between features and removed the 488 features with the highest pairwise correlations. Finally, we removed another 44 features if their minimum or maximum absolute value was greater than 15 (using the default threshold in pycytominer^51^). Hence, we obtained 184 Cell Painting features for 9,876 unique compounds for the dataset comprising public assays. Analogously, for the Broad Institute dataset, we obtained 191 Cell Painting features for 15,272 unique compounds (both datasets are available on Zenodo at https://doi.org/10.5281/zenodo.7589312).

Next, we performed feature selection for the structural features of Morgan fingerprints. For the public assays, we removed 1,891 bits that did not pass a near-zero variance (0.05) threshold since they were considered to have less predictive power. Finally, we obtained Morgan fingerprints of 157 bits for 9,876 unique compounds. Analogously, for the Broad Institute dataset, we obtained Morgan fingerprints of 277 bits for 15,272 unique compounds (both datasets are available on Zenodo at https://doi.org/10.5281/zenodo.7589312).

### Chemical and Morphological Similarity

We next defined the structural similarity score of a compound as the mean Tanimoto similarity of the 5 most similar active compounds. The morphological similarity score of a compound was calculated as the median Pearson correlation of the 5 most positively correlated active compounds.

### Model Training

For each assay, the data was split into training data (80%) and held out test data (20%) using a stratified splitting based on the assay hit call. First, on the training data, we performed 5-fold nested cross-validation keeping aside one of these folds as a test-fold, on the rest of the 4 folds. We trained separate models, as shown in Figure 1c step (1) and step (2), using Morgan fingerprints (157 bits for the public dataset; 277 bits for the Broad Institute dataset) and Cell Painting data (184 features for the public dataset, 191 features for the Broad Institute dataset) respectively for each assay. In this inner fold of the nested-cross validation, we trained separately, Random Forest models on the rest of the 4 folds with Cell Painting and Morgan fingerprints. These models were hyperparameter optimised (with parameter spaces as shown in Supplementary Data 4) using cross-validation with shuffled 5-fold stratified splitting. For hyperparameter optimisation, we used a randomized search on hyperparameters as implemented in scikit-learn 1.0.1^50^. This optimisation method iteratively increases resources to select the best candidates, using the most resources on the candidates that are better at prediction.^52^ The hyperparameter optimised model was used to predict the test fold. To account for threshold balancing of Random Forest predicted probabilities (which is common in an imbalanced prediction problem), we calculated on the 4 folds, the Youden’s J statistic^53^ (J = True Positive Rate – False Positive Rate) to detect an optimal threshold. The threshold for the highest J statistic value was used such that the model would no longer be biased towards one class and give equal weights to sensitivity and specificity without favouring one of them. This optimal threshold was then used for the test-fold predictions, and this was repeated 5 times in total for both models using Morgan fingerprints and Cell Painting features until predictions were obtained for the entire training data in the nested cross-validation manner. As the optimal thresholds for each fold were different, the predicted probability values were scaled using a min-max scaling such that this optimal threshold was adjusted back to 0.50 on the new scale. Further for each test-fold in the cross-validation, as shown in Figure 1c step (3) and step (4), we also calculated the chemical and morphological similarity (as described above in the “Chemical and Morphological Similarity” section) for each compound in this test-fold with respect to the active compounds in the remaining of the 4 folds. This was also repeated 5 times in total until chemical and morphological similarity scores were obtained for the entire training data.

Finally, on the entire training data, two Random Forest models were trained with Cell Painting and Morgan fingerprints with hyperparameter-optimised (in the same way as above using 5-fold crossvalidation). This was used to predict the held-out data, as shown in Figure 1c step (5) (with threshold balancing performed from cross-validated predicted probabilities of the training data). We calculated the chemical and morphological similarity of each compound in the held-out data compared with all active compounds in the training data and these were recorded as the chemical and morphological similarity scores respectively of the particular compound in the held-out dataset as shown in Figure 1c step (6). The predicted probability values were again adjusted using a min-max scaling such that this optimal threshold was 0.50 on the new scale.

### Similarity-based merger model

The similarity-based merger models presented here combined individual scaled predicted probabilities from individual models trained on Cell Painting and Structural data and the morphological and structural similarity of the compounds with respect to active compounds in the training data. In particular, for each assay, we evaluated the similarity-based merger model on the held-out data using information from the training data only to avoid any data or model leakage. We trained a Logistic Regression model (with baseline parameters of L2 penalty, an inverse of regularization strength of 1 and balanced class weights) on the training data which uses the Cell Painting and Morgan fingerprints models’ individual scaled predicted probabilities and the structural and morphological similarity scores (with respect to other folds in the training data itself) as features and the endpoint as the assay hit call of the compound, as shown in Figure 1c step (7). Finally, this logistic equation was used to predict the assay hit call of the held-out compounds (which we henceforth call the similarity-based merger model prediction) and an associated predicted probability (which we henceforth call similarity-based merger model predicted probability), as shown in Figure 1c step (8). There is no leak of any held-out data assay hit call information but only its structural similarity and morphological similarity to the active compounds in the training data, which can be easily calculated for any compound with a known structure.

### Baseline Models

For baseline models, we used two models, namely a soft-voting ensemble^21^ and a hierarchical model^22^. The soft-voting ensemble, as shown in Figure 1a, combines predictions from both the Cell Painting and Morgan fingerprints models using a majority rule on the predicted probabilities. In particular, for each compound, we averaged the re-scaled predicted probabilities of two individual models, thus in effect creating an ensemble with soft-voting. We applied a threshold of 0.50 (since predicted probabilities from individual models were also scaled to the optimal threshold of 0.50 as described above) to obtain the corresponding soft-voting ensemble prediction.

For the hierarchical model, as shown in Figure 1b, we fit a baseline Random Forest classifier (hyperparameter optimised for estimators [100, 300, 400, 500] and class weight balancing using stratified splits and 5 fold cross validations as implemented in scikit-learn^50^) on the scaled predicted probabilities for the entire training data from both individual the Cell Painting and Morgan fingerprints models (obtained from the nested-cross validation). We used this hierarchical model to predict the activity of the held-out test set compounds which gave us the predicted assay hit call (and a corresponding model predicted probability) which we henceforth call the hierarchical model prediction (and a corresponding hierarchical model predicted probability).

### Model evaluation

We evaluated all models (both individual models, soft-voting ensemble, hierarchical and similarity-based merger model) based on precision, sensitivity, F1 scores of the minority class, specificity, balanced accuracy, Matthew’s Correlation Coefficient (MCC) and Area Under Curve-Receiver Operating Characteristic (AUC) scores.

### Statistics and Reproducibility

A detailed description of each analysis’ steps and statistics is contained in the methods section of the paper. Statistical methods were implemented using the pandas Python package.^54^ Machine learning models, hyperparameter optimisation and evaluation metrics were implemented using scikit-learn^50^, a Python-based package. We have released the datasets used in this study which are publicly available at Zenodo (https://doi.org/10.5281/zenodo.7589312). We released the python code for the models which are publicly available on GitHub (https://github.com/srijitseal/Merging_Bioactivity_Predictions_CellPainting_Chemical_Structure_Similarity).

## DECLARATIONS

### ETHICAL APPROVAL

Not applicable.

### AUTHORS’ CONTRIBUTIONS

S Seal designed and performed exploratory data analysis, and implemented, and trained the models. S Seal, H.Y. and S Singh contributed to the design of the models. S Seal, M.A.T and H.Y contributed to analysing the results of the models. S Seal, J.C.P and O.S analysed the biological interpretation of performance. S Seal wrote the manuscript with extensive discussions with O.S. and A.B. who supervised the project. All the authors (S Seal., H.Y, M.A.T, S Singh, J.C.P, O.S and A.B) reviewed, edited, and contributed to discussions on the manuscript.

### Availability of data and materials

We have released the datasets used in this study which are publicly available at Zenodo at https://doi.org/10.5281/zenodo.7589312. We released the python code for the models which are publicly available on GitHub at https://github.com/srijitseal/Merging_Bioactivity_Predictions_CellPainting_Chemical_Structure_Similarity. The protein-protein network (annotated by genes) from the STRING can be accessed at https://version-11-5.string-db.org/cgi/network?networkId=bpH0WmWa1eZm.

Supplementary Data 1: Assay descriptions of the public dataset comprising 88 assays.

Supplementary Data 2: Assay types and readout types of Broad Institute dataset comprising 89 assays.

Supplementary Data 3: Performance of Cell Painting, structural models, soft-voting ensembles, hierarchical models, and the similarity-based merger models over all 177 assays used in this study.

Supplementary Data 4: Hyperparameters considered for optimising Random Forests

## ACKNOWLEDGMENT

The authors are grateful for guidance from Anne Carpenter (Broad Institute of MIT and Harvard) and Juan Caicedo (Broad Institute of MIT and Harvard) which helped improve an earlier version of the manuscript. S Seal would like to acknowledge Steffen Jaensch (Steffen Jaensch) for a discussion which helped correct an important mistake in the earlier version of the manuscript causing an information leak. S Seal would like to thank Chaitanya K. Joshi (University of Cambridge) and Alicia Curth (University of Cambridge) for useful discussions about this work. This work was performed using resources provided by the Cambridge Service for Data Driven Discovery (CSD3) operated by the University of Cambridge Research Computing Service (www.csd3.cam.ac.uk), provided by Dell EMC and Intel using Tier-2 funding from the Engineering and Physical Sciences Research Council (capital grant EP/P020259/1), and DiRAC funding from the Science and Technology Facilities Council (www.dirac.ac.uk). This work was also performed using the Nest compute server (https://www.ch.cam.ac.uk/computing/nest-compute-server) provided by the Yusuf Hamied Department of Chemistry. The TOC Figure and Figure 1 were partly generated from https://bioicons.com/ which uses Servier Medical Art, provided by Servier, licensed under a Creative Commons Attribution 3.0 unported license

